# HIV-1 derived replication intermediates/oligonucleotides induce a type I IFN-dependent immune suppression via STING activation that can be restored by targeting IFNARI

**DOI:** 10.1101/2025.02.17.638698

**Authors:** Cecilia Svanberg, Ravi Prasad Mukku, Özkan Besler, Francis R. Hopkins, Christopher Sjöwall, Sofia Nyström, Esaki M. Shankar, Marie Larsson

## Abstract

The hallmark of HIV-1 infection is the progressive development of multicellular and systemic immune dysfunction, culminating in AIDS. Dendritic cells (DCs) play a pivotal role in HIV dissemination to CD4+ T cells, which are subsequently depleted by the virus leading to HIV disease progression. Type I interferons (IFNs) are critical for host defense during acute infection but contribute to chronic immune activation during the later stages of HIV disease. This persistent activation leads to immune cell exhaustion. HIV-1 can activate type I IFN responses via several pathways, including the STING pathway, which is activated by e.g., virus-derived oligonucleotides. Here, we investigated the underlying mechanisms creating HIV-1-mediated immune dysfunction and role of type I IFNs using a DC and T cell co-culture model. HIV-1 exposure in the DC-T cell co-culture promoted the expansion of suppressive T cells with diminished proliferation and effector functions. The impairment required type I IFNs and subsequent IFN-α/β receptor signaling, initiated by HIV-derived ssDNA activation of IFI16/cGAS followed by STING signaling in the DCs. Targeting IFNAR1 with Anifrolumab restored the immune functions of both DCs and T cells, as well as T cell proliferation and T cell effector functions, including their secretion of IL-2, IFN-γ, and granzyme B. Our findings support that the immune impairments existing in untreated or antiretroviral therapy (ART) treated HIV-infected individuals are mediated, if not fully in part by type I IFN’s negative effect on DC and T cells. Therapeutics targeting IFN-α/β receptors, such as Anifrolumab, hold potential as combination treatment alongside ART, to achieve a more complete immune restoration and contribute to improved quality of life among people living with HIV.

## Introduction

Over three decades since its discovery, human immunodeficiency virus type 1 (HIV) infection continues to be a major global public health challenge. As of 2022, more than 38 million individuals live with HIV, with ∼1.3 million new infections reported (UNAIDS, 2022). Encouragingly, the incidence of new infections is declining, attributed to that over 75% of the people living with HIV-1 have access to antiretroviral therapy (ART). The therapy provides effective control of viral replication and limits the negative effect HIV exerts on the immune system and general health (WHO, 2022).

Dendritic cells (DCs) are pivotal players in the initiation and regulation of immune responses, as the functionality of these cells in the lymphoid tissues determines the quality of the ensuing immune response^1,2,3^. DCs are among the first immune cells targeted by HIV upon infection of the genital mucosa, where they reside within the mucosal epithelium^3^. The mucosal DCs are less permissive to infection than CD4+ T cells, but relatively more permissive than blood DCs^4,5^. Still, the DCs exhibit remarkable efficiency in facilitating HIV infection while in cellular contact by transferring virions to the interacting T cells even in the presence of ART^6,7^. Of note, in HIV-infected individuals, DCs in lymphoid tissues can serve as a source of latently infected cells despite years of suppressive ART. These lymphoid located myeloid DCs also express programmed death-1 ligand 1 (PD-L1) in combination with an activated phenotype, which might impair the activation of HIV-specific T cells^8,9^.

Chronic HIV-1 infection is characterized by profound immune dysfunction and systemic immune activation. The immune activation is driven by several factors, and prolonged and elevated type I IFN levels are detrimental to immune responses during chronic HIV infection, as previously demonstrated both in HIV and simian immunodeficiency virus (SIV) infections, where type I IFN signaling contributes to disease progression^10,11^. Many immune cells, including T cell and DC subsets, are functionally impaired in HIV infected, with altered cytokine profiles, activation status, and elevated levels of immune checkpoint molecules, caused by direct HIV effects or by virus-induced bystander effects^12–15^. Importantly, the impaired T cell function in HIV-infected individuals with or without ART can be partially restored through *ex vivo* blockade of immune checkpoint molecules^16^. Anifrolumab, a fully human immunoglobulin G1κ monoclonal antibody that binds to the type I interferon receptor subunit 1 and inhibits signaling by all type I interferons, is approved for moderate-to-severe systemic lupus erythematosus since 2021^17^.

We have demonstrated that during the interaction between mature DCs and T cells in the presence of HIV, the virus induces impaired responses with expression of e.g., PDL1 and galectin-9 on DCs, and programmed death-1 (PD-1) and CTLA4 on T cells^18–20^, which is in line with findings in chronic HIV infection^14,21^. In addition, we demonstrated that the transcriptome profile of HIV-exposed DCs engaged in crosstalk with T cells exhibit a robust prolonged type I IFN signature^20^.

During retroviral and DNA virus infections, oligonucleotides are generated at various stages of the viral life cycle and in the context of HIV an array of replication intermediates such as ssDNA, dsDNA, RNA:DNA hybrids be found in the cytosol^22,23^. These viral nucleic acids are recognized by innate immune sensors such as retinoic acid-inducible gene I (RIG-I), which detects RNA:DNA hybrids^24^, and cyclic GMP-AMP synthase (cGAS) and IFN-inducible protein 16 (IFI16), that trigger type I IFN responses via the stimulator of interferon genes (STING) pathway^22,23,25^. In DCs, the intracellular detection of HIV has been deemed to be dependent on cGAS ^26,27^. In addition to cytosolic sensing, in endosomal compartments containing TLR8 in myeloid DCs and TLR7 and TLR9 in plasmacytoid DCs, the degraded virus is detected even in the absence of productive viral replication^28,29^. The activation of the STING pathway and subsequent expression of type I IFNs has been associated with the induction of immunoregulatory factors such as indoleamine 2, 3 dioxygenase (IDO), PD1, PDL1, and PDL2, contributing to the suppression of T cell proliferation and a general immunosuppressive microenvironment^30,31^.

To investigate the role of type I IFN in suppressive responses induced by HIV-exposed DCs, we examined the HIV replication intermediates driving type I IFN activation and the signaling pathways involved in immune modulation/impairment. To achieve this, we utilized a HIV exposed or unexposed mature DC-T cell model of immune activation, mirroring immune response initiation occurring in lymphoid tissue in an HIV-infected individual.

Our results show that HIV suppressed the DCs and T cells, independent of the HIV-1 M strains used (BaL vs THRO), and that there was a strong and prolonged type I IFN response in HIV-exposed co-cultures. HIV replication intermediates/oligonucleotides participated in the HIV-induced suppression, with ssDNA being the most potent and the IFI16/cGAS/STING signaling pathways the main signaling pathways contributing to the induction of suppressive responses. The negative effects on DCs and T cells in the co-culture were reverted by addition of the anti-IFNAR1 antibody Anifrolumab, blocking the IFNaR1 signaling pathway. Therapies targeting IFNα/β receptors, such as Anifrolumab, hold potential as combination treatment alongside ART to accomplish complete immune restoration and thereby improve health in people living with HIV.

## Introduction

### Cell culture media and reagents

Cell culture was conducted in RPMI1640 medium (ThermoFisher, Stockholm, Sweden) supplemented with Gentamicin (ThermoFisher), HEPES (ThermoFisher), and either 5% pooled human serum (PHS) (Innovative Research, Novi, USA) or 1% single donor human plasma. The ssDNA, dsDNA, various DNA loops, and DNA:RNA hybrids (Invivogen, Toulouse, France), derived from sequences described by Jakobsen et al.^23^ **(Supplementary Table 1)**, were diluted in pH adjusted potassium chloride buffer and added to the cells in amounts ranging from 2 to 16 µg. Recombinant human granulocyte-macrophage colony-stimulating factor (rhGM-CSF) (Prepotech, London, UK) at 100 IU/mL and recombinant human interleukin-4 (rhIL-4) (Prepotech) at 300U/mL were utilized to differentiate monocytes into DCs *in vitro*. To investigate the STING signaling pathway, the following reagents and concentrations were employed: cGAMP (10µg/mL, Invivogen), MSAD2 (33µM, MedChemExpress Europe, Sollentuna, Sweden), ADS100 (10µM, MedChemExpress), SN011 (1 µM, MedChemExpress), 5′pppRNA (1µg/mL, Invivogen), RIG012 (1.25 µM, Axon Medchem, Groningen, Netherlands), and VACV70 (10µg/mL, Invivogen). Additional chemicals and compounds used included CL075 (1µg/mL, Invivogen), recombinant p24 (1µg/mL, Invivogen), and IFNAR blockade using Anifrolumab at 20 µg/mL, as a kind gift from AstraZeneca.

### Dendritic cell propagation

Whole blood or leukocyte preparations from healthy volunteers were processed using a Ficoll-Paque™ (Amersham Pharmacia, Piscataway, NJ, USA) density gradient. The PBMCs were subjected to sequential centrifugation at 1800, 1500, 1100, and 900 RPM for 10 minutes each at 4°C. After centrifugation, the cells were counted and plated at a density of 40 million cells per plate in 10 mL of 5% PHS. The plates were incubated at 37°C for 1–2h to allow monocyte adherence. Subsequently, non-adherent cells were removed, and the plates were washed. The remaining adherent monocytes were cultured in 1% plasma medium and differentiated into monocyte-derived DCs by adding GM-CSF and IL-4 every other day for 5 days. On day 5, phenotype was investigated by flow cytometry to exclude undifferentiated cells by CD14 expression and spontaneously activated cells by CD83 expression. The maximum cut-off for both markers was set to ∼15%.

### Maturation of dendritic cells

Following the cut-off selection for viable cells, moDCs were harvested, counted, and re-seeded in 10 mL of the same conditioned culture medium used during differentiation, at a density of 4 million cells per plate. The TLR3 ligand poly I:C (3ng/ml: Invivogen) was then added to the cultures, and the moDCs were incubated at 37°C for 24h to promote cell maturation.

### HIV infection and oligo transfection of moDCs

After 24h of maturation CCR5-tropic HIV BaL 4238 or HIV THRO A66 4393 was added in a concentration of 750 ng/mL and DCs were incubated another 24h. For intracellular addition of nucleotides, proteins, and antagonists/agonists the N-[1-(2,3-dioleoyloxy)propyl]-N,N,N-trimethylammonium methylsulfate (DOTAP) (Merck, Darmstadt, Germany) liposomal transfection system was used. An amount of 30µl of DOTAP was mixed with 70µl of HBSS buffer (0.9% NaCl+ 10mM HEPES) and thereafter the oligos or proteins was added in indicated concentrations diluted in 50µl of HBSS. The combined mixture was then incubated for 30 min at room temperature. The whole volume was added dropwise to 0.5-2 million resuspended mature DCs in 100 µl of culture media. Thereafter the cells were incubated for 2h at 37°C, spun down and washed to remove the DOTAP mixture and resuspended in conditioned media at a concentration of 1 million cells/mL.

### Naïve T cell enrichment and set up of allogenic co-cultures

Nonadherent cells from the DC propagation were counted and incubated with 20 µl of buffer and 10µl of each MACS antibody (anti-CD14, CD56, CD19, and CD45R0) (Miltenyi, Stockholm, Sweden) per 10 million cells for 15 min at 4°C. The labelled cells were washed and then passed through an LD-column attached to a magnet for the negative selection of bulk naïve T cells. After collection of total effluent, the naïve T cells were counted and resuspended to 1 million cells/mL in 5% PHS.

### DC-T cell co-culture

The mature DCs exposed to mock, HIV, transfected, or exposed to agonist/antagonists were harvested and resuspended in 5% PHS. The different DC groups were divided, with one portion frozen down in freezing media and stored at −80°C and the other portion plated in a flat bottomed 96-well cell culture plate. Naïve T cells were added to the different DC groups at a ratio of 1:10 and the DC-T cell co-culture incubated in a CO_2_ incubator at 37°C for 7 days. After 7 days, the DC-T cell co-cultures were re-stimulated with the corresponding DC group and left overnight until harvest.

### ELISPOT and ELISA measurements

For ELISA measurements of type I IFN released into the supernatant a PAN IFNα ELISA kit (Mabtech, Stockholm, Sweden) and an IFNβ ELISA construction kit (Antigenix America inc, New York, NY, USA) was used. Supernatants harvested on day 8 were either diluted 1:5 or used neat. For the ELISA, plates were coated with (1µg/mL for IFNβ and 4µg/mL for IFNα) capture antibody overnight, washed and blocked in 0.05% Tween 20 and 0.1% BSA for 60 min. Samples and standard were added to the plate after washing and incubated for 2h at RT. The biotin tracer antibody was thereafter added in a concentration of 0.5-1µg/mL and incubated for 1h at RT. Streptavidin-HRP was added in 1:1000 dilution for 30 min at RT before addition of TMB substrate. The reaction was terminated after 15 min by the addition of 0.2M H_2_SO_4_ and the plates read at 450 nm using a SpectraMax iD3 microplate reader (Molecular devices, San Jose, CA, USA). For ELISPOT assays cells from the coculture were counted on day 7 and replated onto pre-coated plates for human granzyme B, and IFNγ (Mabtech). For IL2 PVDF plates were coated in house by addition of 10µg/mL of capture antibodies (Mabtech). Thereafter DCs were added at 1:10 ratio to each well and plates were incubated at 37°C overnight. Spots were detected using biotin antibodies (0.25-0.5µg/mL) and streptavidin-HRP (1:1000 dilution). TMB was used to develop the color and the reaction stopped by washing with deionized H_2_O and spots were counted manually in a light microscope.

### 3H-thymidine incorporation proliferation assay

To measure T cell proliferation in the DC-T cell co-culture during restimulation, 2µCi/µl of ^3^H-thymidine (PerkinElmer, Waltham, MA, USA) was added following the supplement of the DCs. Wells containing RPMI and no cells were used for background counts. The amount of incorporated ^3^H-thymidine was measured using liquid scintillation and counted using a micro-β-counter (PerkinElmer).

### Statistics

For data normalized to mock equals 1, a Kruskall-Wallis test followed by Dunn’s multiple comparisons test was performed. In the case of qPCR, where the sum of values from each donor was set to 100%, a One-way ANOVA with Tukey’s post-hoc test was performed. All tests were done using GraphPad Prism version 9.2.1.

## Results

### HIV-1 BaL and HIV-1 THRO had similar suppressive effects on the DC priming of naïve T cells

To investigate whether the presence of HIV-1 during the DC priming and restimulation of T cells affected the ensuing immune response we co-cultured naïve T cells with DCs exposed to different strains of CCR5-tropic HIV-1, either HIV-1 BaL a late-stage derived lab adapted strain or HIV-1 THRO a transmitted/founder strain. Our analysis revealed a significant suppression of T cell proliferation **(Figure 1A)**, and IL-2 production **(Figure 1B)** for both HIV-1 strains but no significant differences in outcomes between the two strains. There were elevated gene transcripts of DC and T cell immunomodulatory factors **(Figure 1C)** for both HIV-1 strains, but no significant differences observed in outcomes besides significantly higher PDL1 in HIV-1 THRO compared to HIV-1 BaL. Next, we explored induction of type I IFN responses and ISGs in the DC-T cell co-cultures **(Figure 1D)** and discovered an upregulation of IFNα4 and several ISGs by both HIV-1 strains.

**Figure 1.**
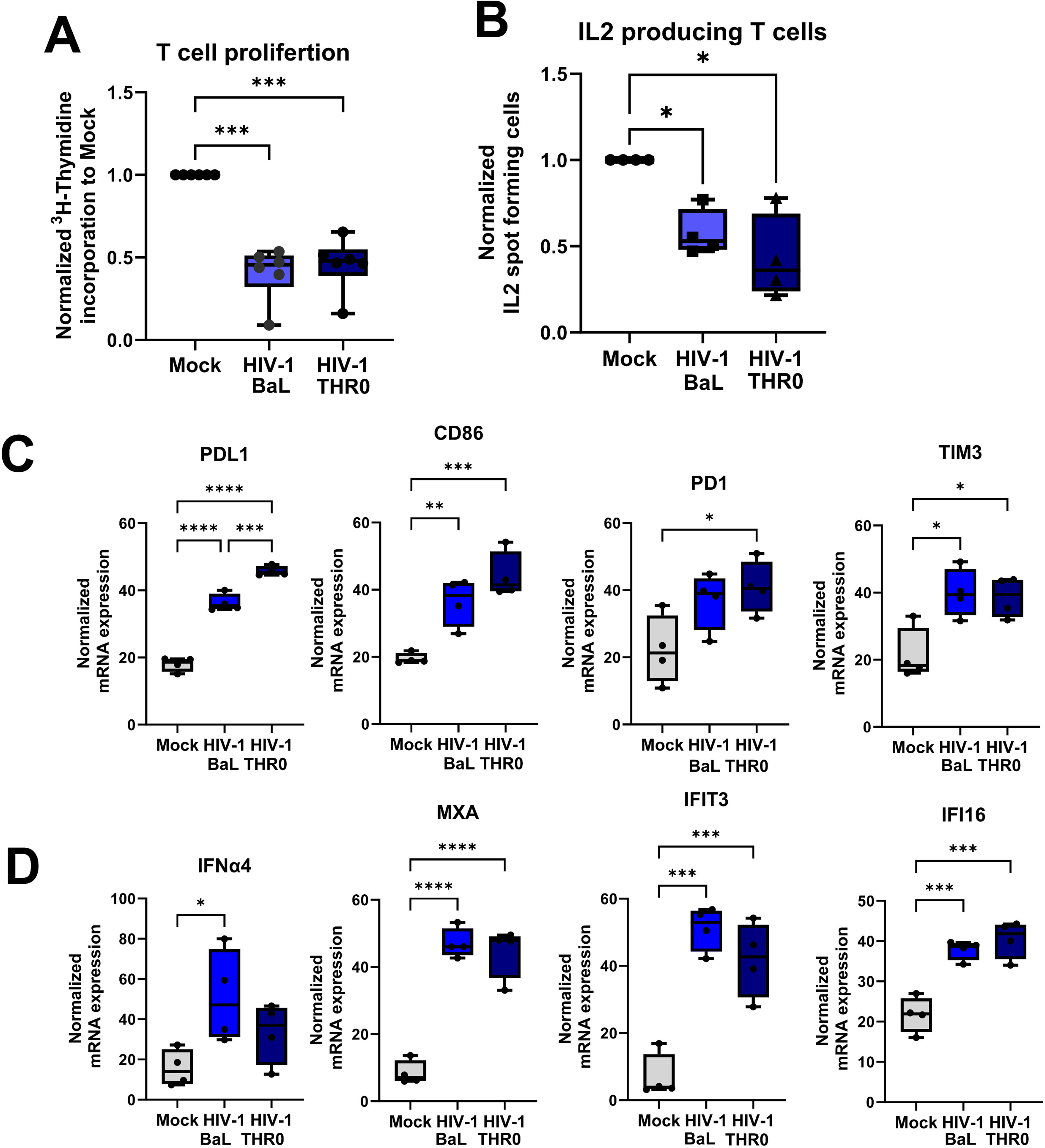
Both chronic and founder HIV-1 strains suppressed dendritic cells ability to prime naïve T cells. HIV-1 BaL a late-stage derived lab adapted strain or HIV-1 THRO a transmitted/founder strain exposed mature DCs were co-cultured with naïve T cells in the ratio of 1:10. The DC-T cell co-culture was re-stimulated on day 7 with the same DCs as the initial stimulation. One day after restimulation, the T proliferation was assessed via ^3^H thymidine incorporation **(A)** and functionality by enumerating lL2-producing cells by ELISPOT assays **(B)**. mRNA transcript of PDL1, CD86, PD1, and TIM3, and **(C)** IFNα4, MXA, IFI13, and IFI16 **(D)** by qPCR. Statistical significance was determined using the ANOVA. * =p-value <0.05, ** = p-value <0.01, *** = p-value <0.001**** = p-value <0.0001.

### Pathways inducing type I IFNs and type IFN responses were enriched and predicted to be activated when HIV was present in DC-T cell co-culture and exogenous type I IFN resulted in suppressed T cell responses similar to HIV

To establish pathways involved in HIV suppression, we examined top regulator networks in an RNA sequencing data set from a previous study on HIV exposed DCs from DC-T cell co-culture^20^. Therein, we found a strong activation of intracellular sensors involved in type I IFN signaling and responses, (e.g., sGAS, STING, and MAVS), IFN regulatory transcription factors (e.g., IRF3, IRF9, and STAT1), type I IFNs, and IFNAR1 and IFNAR2 **(Figure 2A).** Therefore, to explore if type I IFN played a key role in the induced T cell suppression in our system, we added recombinant type I IFNα2 protein to the DC-T cell co-cultures. The IFNα2 induced suppressed T cell responses were similar to those induced by HIV in the DC-T cell co-culture **(Figure 2B),** even at concentrations similar to the levels detected in the DC-T cell co-culture exposed to HIV-1 **(Figure 1D and Figure 2B)**.

**Figure 2.**
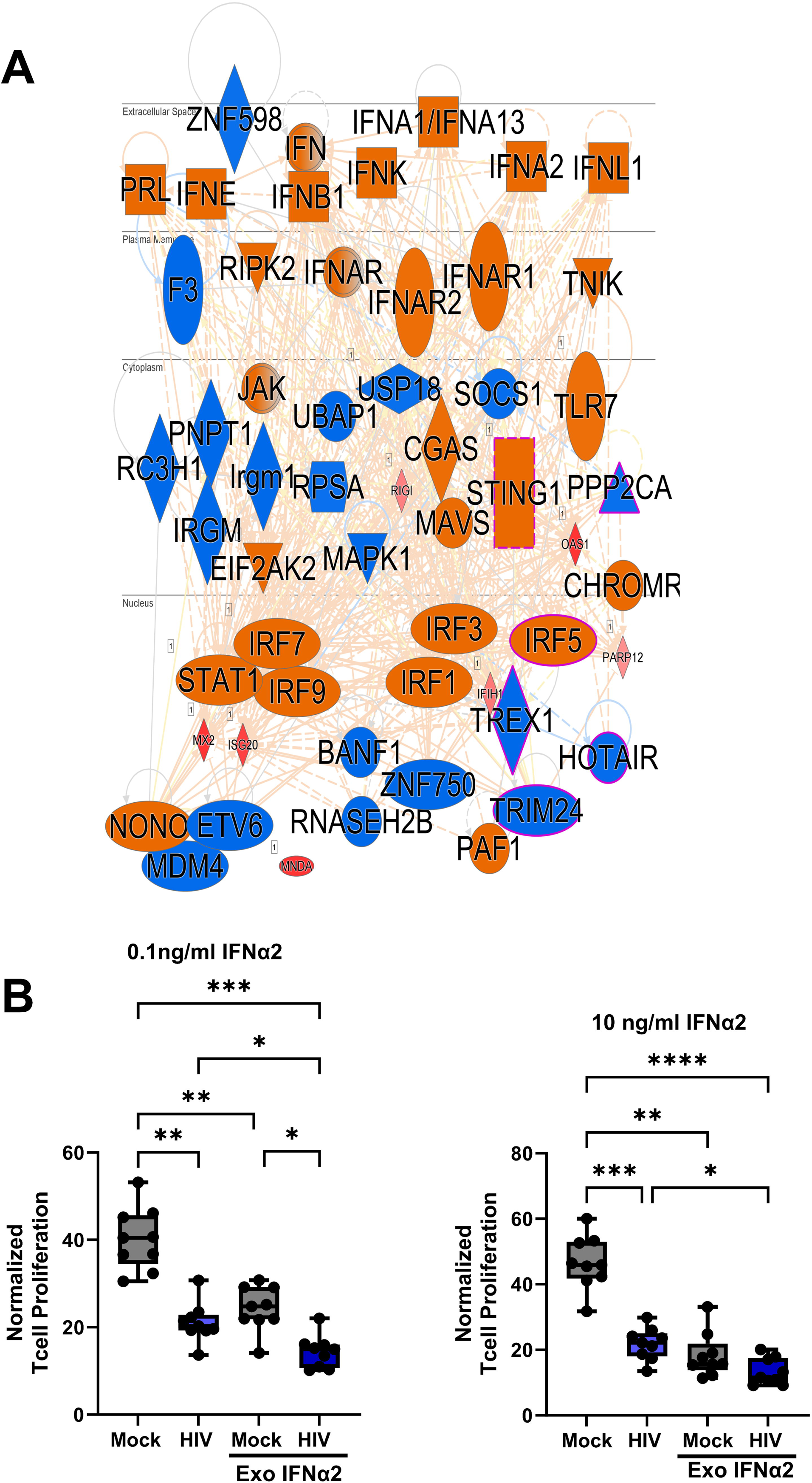
Type I IFN inducing pathways were enriched and predicted to be activated when HIV was present in DC-T cell co-culture and exogenous type I IFN-induced suppressed T cell responses similar to HIV. Transcriptomic data from our published RNA seq^20^, where DCs purified from the DC-T cell co-culture after one day of restimulation, were assessed for top regulator network **(A).** HIV-1 BaL exposed mature DCs were co-cultured with naïve T cells in the presence or absence of exogenous human recombinant IFNα4. The DC-T cell co-culture was re-stimulated on day 7 with the same DCs as the initial stimulation and the IFNα2 replenished. One day after restimulation, the T proliferation was assessed via ^3^H thymidine incorporation **(B)**. mRNA transcript of PDL1, and PD1 **(C)** by qPCR. Statistical significance was determined using the ANOVA. * =p-value <0.05, ** = p-value <0.01, *** = p-value <0.001**** = p-value <0.0001.

### HIV-derived oligonucleotide ssDNA participated in the induction of the suppressive effects seen by HIV in the DC-T cell co-culture

We explored which of the HIV-1 derived oligonucleotides or proteins produced during the viral life cycle that could be involved in the HIV-induced immune impairment of the DCs and T cells during priming. The ssDNA, dsDNA, DNA:RNA hybrid, many loops ssDNA, and p24 were delivered intracellularly into the DCs by DOTAP transfection **(Figure 3)**. We found that ssDNA had the most potent suppressive effects on the T cell proliferation and that dsDNA also had negative effects on T cell activation but was more variable **(Figure 3A).** The RNA:DNA hybrid had significant suppressive effect but much less effect than seen for the ssDNA **(Figure 3A-B)**. To explore the effect of HIV-1 genome, i.e., ssRNA, we delivered a custom made ssRNA **(Supplementary Table 1)** and the synthetic ssRNA40 intracellularly to the DCs and found that they had no or negligible effect on the T cell proliferation (**Figure 3C**). The HIV-1 p24, and many loops ssDNA, had no effects on the T cell proliferation **(Figure 3B and D)**. To ensure that the transfection did not affect the DC-T cell priming/activation, we treated HIV-exposed or unexposed DCs with DOTAP and noticed no significant effects **(Supplementary Figure 1)**. The HIV-1 derived oligonucleotides known to induce STING signaling^23^ had a negative effect on T cell proliferation and among them ssDNA had the most potent and consistent effects and was selected for further studies of the mechanism behind the HIV-induced impairment.

**Figure 3.**
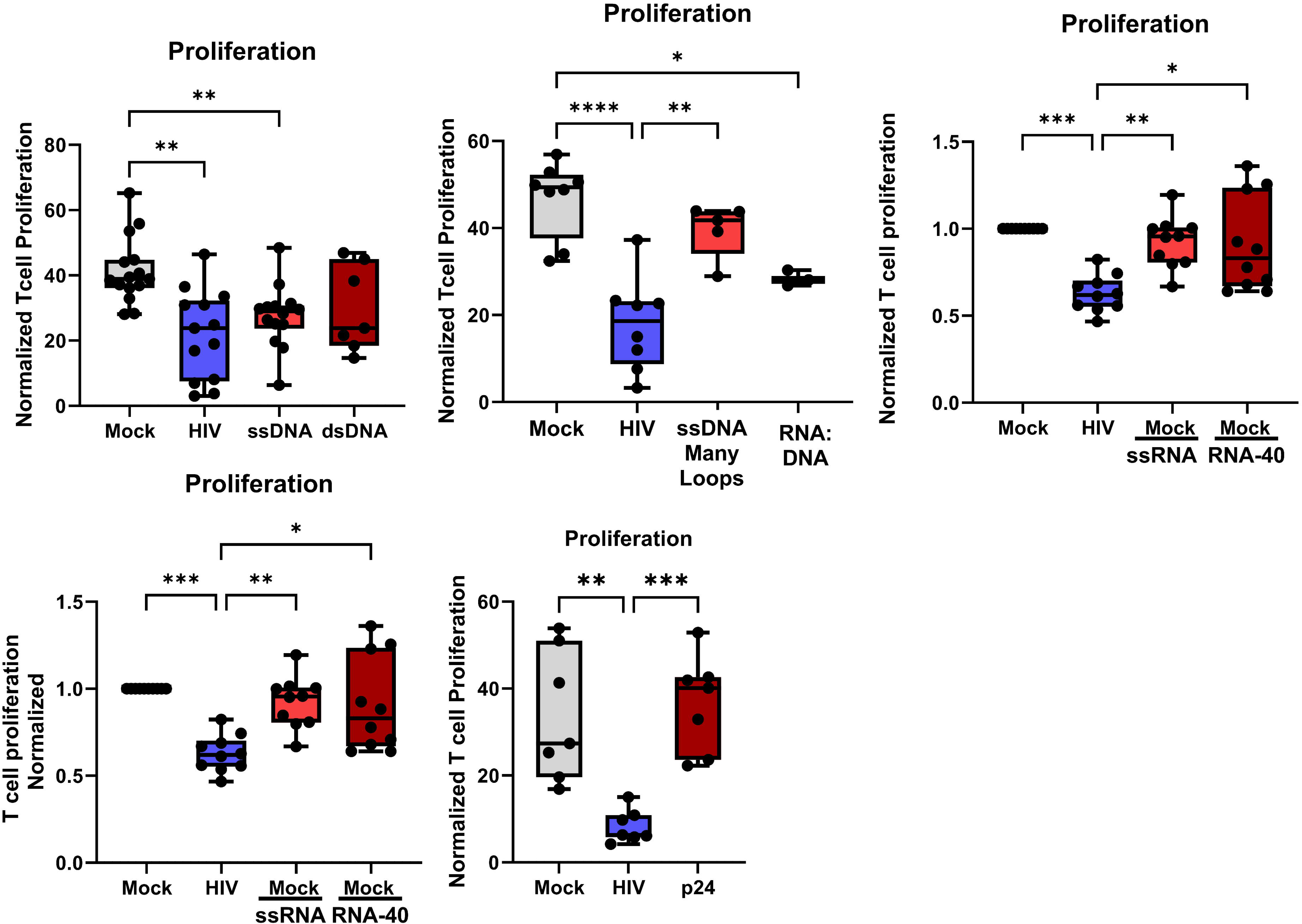
HIV-1 derived replication intermediate ssDNA participated in the induction of the suppressive effects seen by HIV in the DC-T cell co-culture. Mature DCs were exposed to HIV-1 BaL or transfected with the ssDNA, dsDNA, DNA:RNA hybrid, many loops ssDNA, or p24 using DOTAP to deliver these HIV components into the cytosol. The different DC groups were co-cultured with naïve T cells (1:10) and the DC-T cell co-culture was re-stimulated on day 7 with the same DCs as the initial stimulation. One day after restimulation, the T proliferation was assessed via ^3^H thymidine incorporation **(A-D).** Statistical significance was determined using the ANOVA. * =p-value <0.05, ** = p-value <0.01, *** = p-value <0.001**** = p-value <0.0001.

### HIV and ssDNA induce type I IFN production and ISG in DC-T cell co-cultures

HIV-1, ssDNA, and dsDNA, induced mRNA transcript of IFNβ, IFNα and ISGs such as IFI16 (**Figure 4A-B**). Of note, even if HIV-1, ssDNA, and dsDNA gave rise to type I IFN and ISGs on mRNA levels, only the complete virion induced detectable amounts of IFNα and IFNβ on protein level measured by ELISA (**Figure 4C**). HIV-1 induced IFNα and IFNβ on day 3 after co-culture setup and the production persisted throughout the DC-T cell co-culture (**Figure 4C)**.

**Figure 4:**
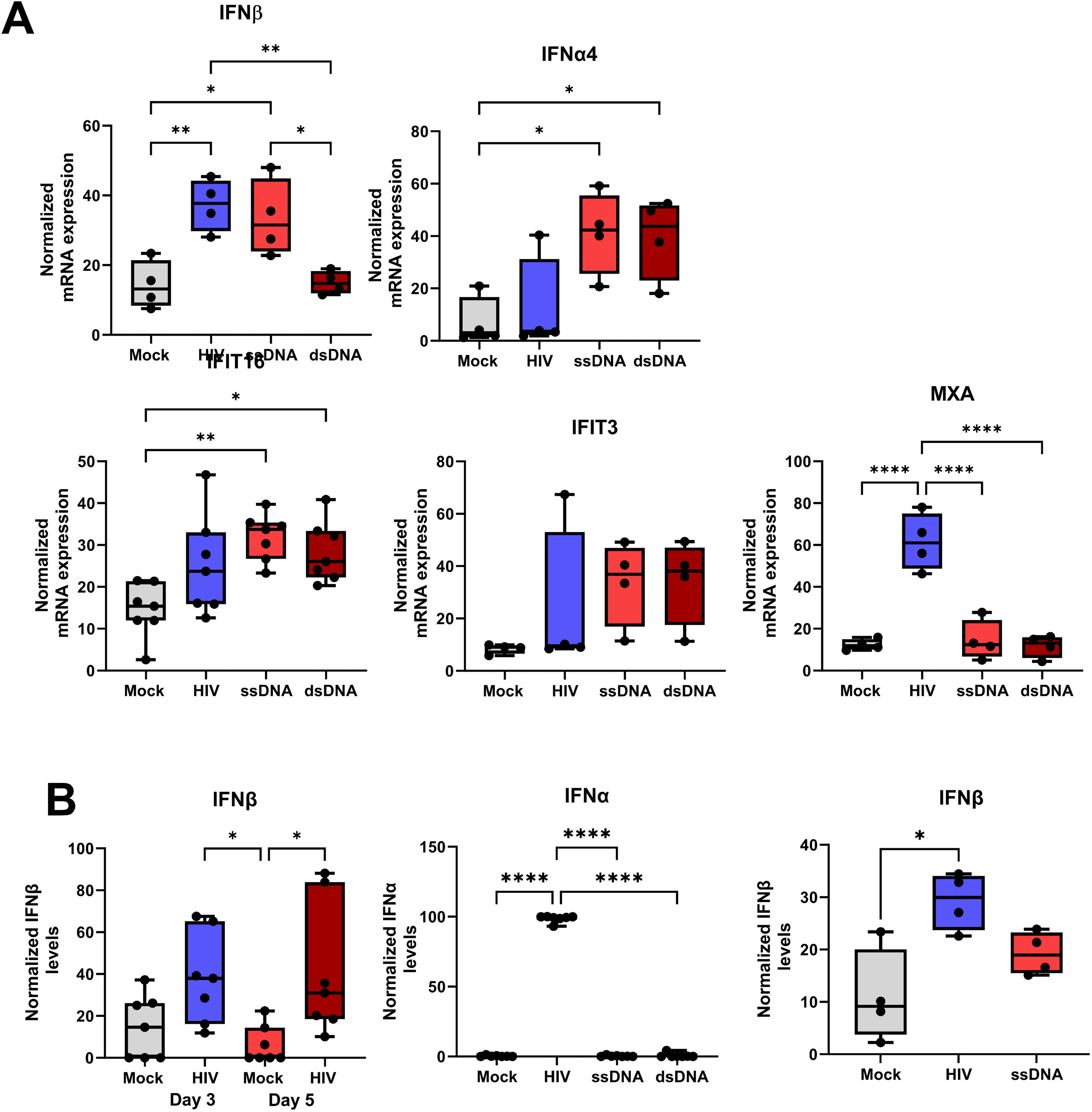
HIV and ssDNA induce type I IFN production and ISG in DC-T cell co-cultures. Mature DCs were exposed to HIV-1 BaL or transfected with the ssDNA, dsDNA, DNA:RNA hybrid, many loops ssDNA, or p24 using DOTAP to deliver these HIV components into the cytosol. The different DC groups were co-cultured with naïve T cells (1:10) and the DC-T cell co-culture was re-stimulated on day 7 with the same DCs as at the initial stimulation. mRNA transcript of **(A)** IFNα4, MXA, IFI13, and IFI16 by qPCR and **(B)** Protein levels of type I IFNs, IFNα4 and IFNβ were measured by ELISA. ANOVA. * =p-value <0.05, ** = p-value <0.01, *** = p-value <0.001, **** = p-value <0.0001.

### STING agonists suppressed T cell proliferation and increased PD-L1 expression on the cells in the co-culture

Activation of STING by DNA oligonucleotides is an indirect event involving the activation of DNA sensors e.g. cGAS, and IFI16, and followed by downstream STING activation^30^. To explore the involvement of the STING signaling pathway in the HIV-induced suppression, we targeted the STING protein directly by agonists, i.e., cyclic guanosine monophosphate adenosine monophosphate (cGAMP) produced when cGAS is bound to a DNA ligand^25^, a stabilized 2’3’cyclic-di-AMP analog ADUS100 ^32^, and MSA2^32^. Additionally, we tested the antagonist SN-011 which competes with cyclic dinucleotide for the binding pocket on STING^33^. When activating STING using extracellular or intracellular delivered cGAMP we found some decreased proliferation with the intracellular but not extracellular delivered cGAMP **(Figure 4A)** and for the agonists ADU100 and MSAD2 we found an even stronger suppressive effect on the T cell proliferation **(Figure 4B-C)**. The antagonist SN-011 treated cells gave the same profile as untreated, i.e., it failed to stop/rescue the negative effect exerted by HIV-1 on the T cell proliferation in the DC-T cell co-culture **(Figure 4D)**. Exploring the effect on type I IFNs and ISGs we found induction of IFNβ, MXA, and PDL1 by STING agonists, especially MSA2 **(Figure 4E-G)**. STING activation in T cells has been shown to lead to impaired T cell proliferation and in our hands STING activation gave rise to T cell suppression, which could be type I IFN conditioning of the T cells interacting with the DCs.

### RIG activation played no role in the HIV induced immune suppression in the DC T cell coculture

To further investigate sensors upstream of STING that could be involved in the activation of type I IFNs and in the HIV-induced immune suppression, we explored the effect of the RIG antagonist RIG012^34^. Intracellular inhibition of RIG in the DCs in the DC-T cell co-culture did not affect the T cell suppression induced by HIV-1 **(Figure 5A).** Furthermore, RIG012 inhibition of RIG had no effect on type I IFN or ISGs **(Figure 5B**), and no or insignificant effect on co-inhibitory molecules **(Figure 5C)**. These findings help to exclude the involvement of RIG in HIV-induced immune impairment.

**Figure 5:**
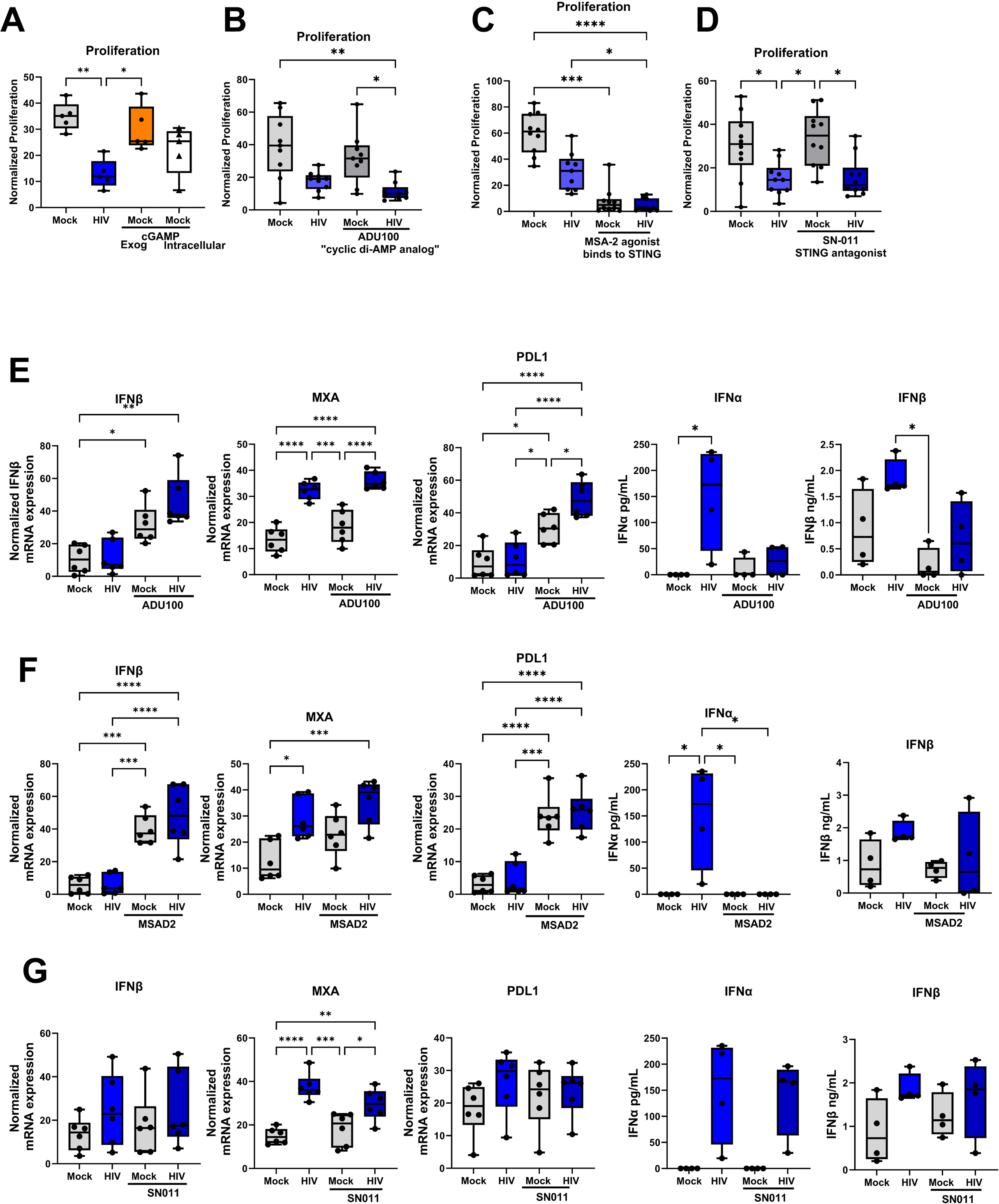
STING agonists suppressed T cell proliferation and increased PDL1 expression of the cells in the co-culture. Mature DCs were unexposed or exposed to HIV-1 BaL, or transfected with STING agonists cGAMP, and cocultured with naïve T cells. Additionally, during coculture the cells were exposed to, stabilized 2’3’cyclic-di-AMP analog ADU100, and MSA-2, and STING antagonist SN-011. Cocultures were restimulated on day 7 by the addition of the same DCs as during setup. One day after re-stimulation, the T proliferation was measured by ^3^H thymidine incorporation **(A-D).** mRNA transcript levels of IFN-β, MXA, and PDL1 in ADU100 **(E)**, MSA2 (F), and SN-011 **(G)** treated DC-T cell co-culture assessed by qPCR. Protein levels of IFNα and IFNβ in ADU100 **(E),** MSA2 **(F)**, and SN-011 **(G)** treated DC-T cell co-culture assessed by ELISA. Statistical significance was determined using the ANOVA. * =p-value <0.05, ** = p-value <0.01, *** = p-value <0.001, **** = p-value <0.0001.

### IFI16 activation suppresses T cell proliferation but is not enough to induce co-inhibitory molecules

To further investigate the upstream sensors of STING involved in HIV-induced immune suppression, we added the IFI16 agonist VACV-70, a dsDNA derived from vaccina virus^23^. The activation of IFI16 sensor suppressed proliferation to similar levels as HIV-1 (**Figure 6A**). Surprisingly, the IFI16 agonist had no or little enhancing effects on the levels of coinhibitory molecules or MXA, compared to HIV in DCs or T cells in the co-cultures (**Figure 6B-C**), whereas the effects on the IFNα, IFI16, and STING was similar to HIV **(Figure 6C)**. This suggests that some of the negative effects induced by HIV can be replicated by the VACV-70 IFI16 activation and indicate that this pathway could be a major contributor to the HIV induced impairment.

**Figure 6:**
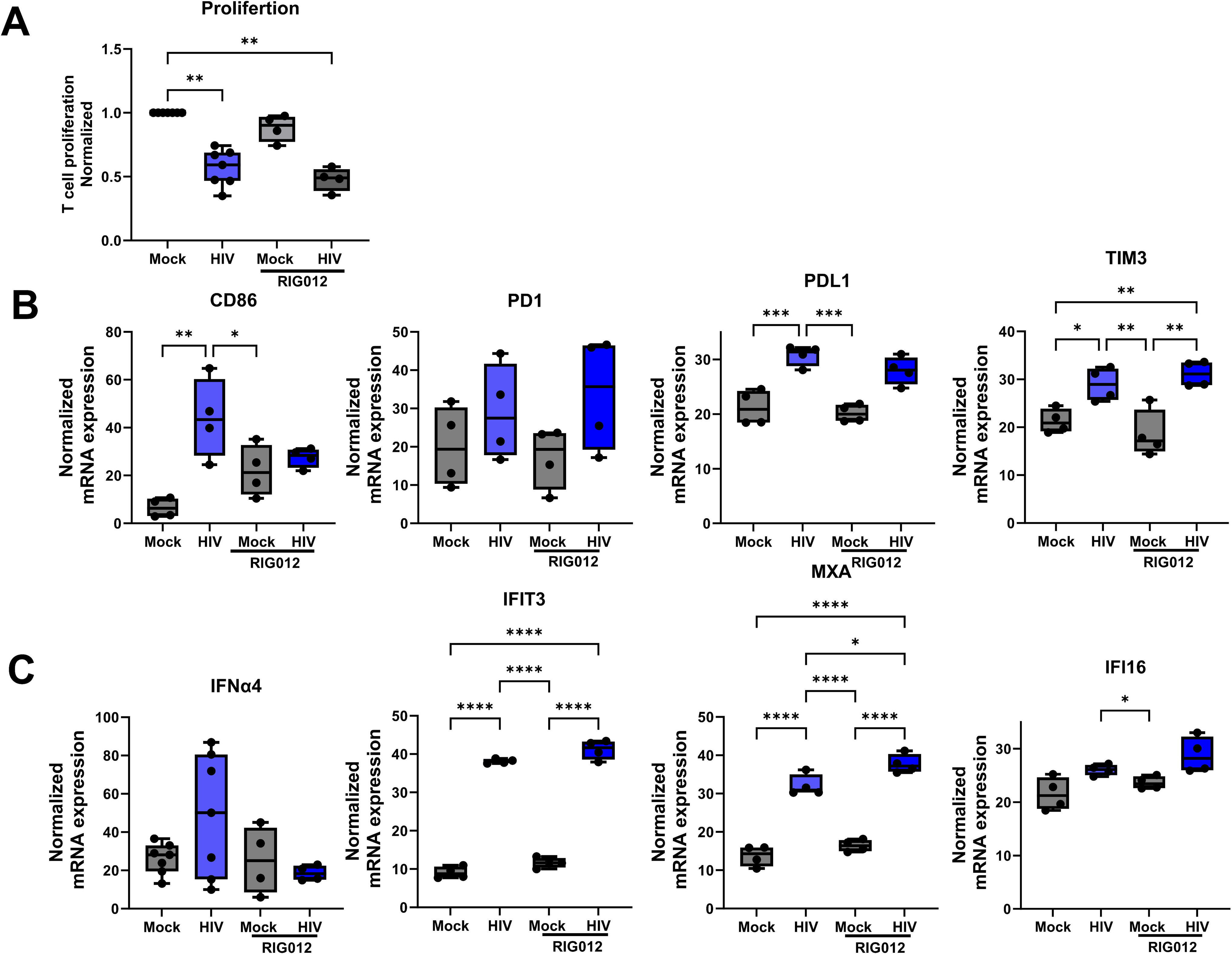
RIG activation played no role in HIV-induced immune suppression in the DC-T cell co-culture. Mature DCs were exposed to HIV-1 BaL or DOTAP transfected with RIG agonist RIG012 to achieve delivery into the cytosol. The different DC groups were co-cultured with naïve T cells (1:10) and the DC-T cell co-culture was re-stimulated on day 7 with the same DCs as the initial stimulation. One day after re-stimulation, the T proliferation measured via ^3^H thymidine incorporation **(A)**, mRNA transcript of CD86, PD1, PDL1, and TIM3 **(B),** and IFNα4, MXA, IFI13, and IFI16 **(C)** by qPCR. Statistical significance was determined using the ANOVA. * =p-value <0.05, ** = p-value <0.01, *** = p-value <0.001**** = p-value <0.0001.

### Blocking of the type I IFN receptor restores DC-T cell function and phenotype

Since we found that the type I IFN response is the major reason for HIV-induced immune suppression, a block of the IFNAR could yield deeper understanding on how this occurs. Blocking of the receptor using Anifrolumab, an anti-IFNAR1 antibody, restored all suppressive events induced by HIV-1, the proliferation and functionality as measured by IFN-α, IL-2 and granzyme B production by T cells (**Figure 7A**), the expression levels of immunomodulatory molecules (**Figure 7B**) and abolished the type I IFN response (**Figure 7C**). This clearly demonstrated the immune suppressive autocrine effect that type I IFNs have via activation of the IFNAR signaling cascade.

**Figure 7:**
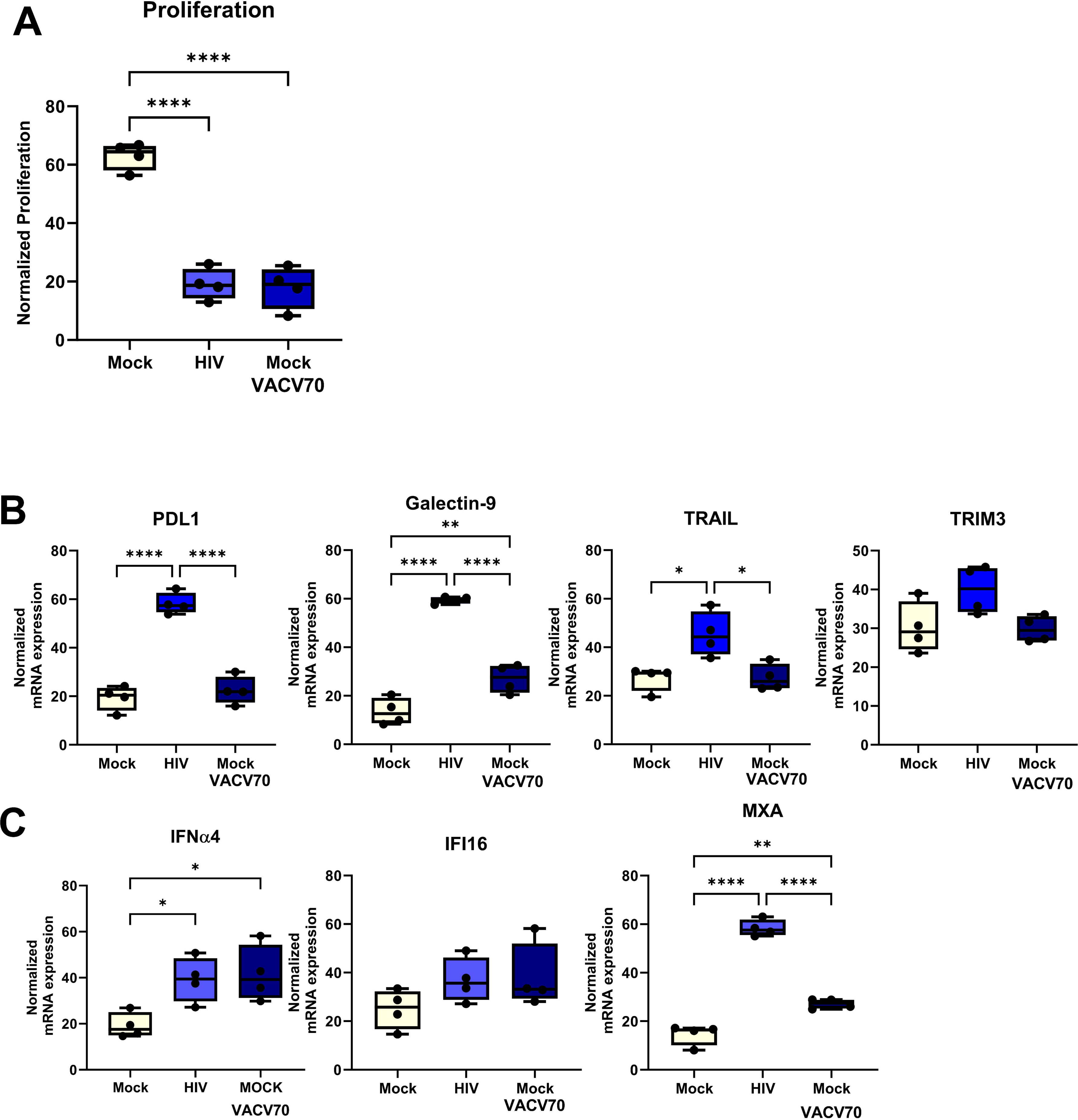
IFI16 activation suppresses T cell proliferation but is not enough to induce co-inhibitory molecules. Mature DCs were exposed to HIV-1 BaL or DOTAP transfected with IFI16 agonist dsDNA VACV-60 to deliver the VACV-60 into the cytosol. The different DC groups were co-cultured with naïve T cells (1:10) and the DC-T cell co-culture was re-stimulated on day 7 with the same DCs as the initial stimulation. One day after re-stimulation, the T proliferation measured by ^3^H thymidine incorporation **(A).** mRNA transcripts of PDL1, Galactin 9, TRAIL, and TRIM3 **(B),** and of IFNα4, IFI16, and MXA **(C)** were assessed by qPCR. Statistical significance was determined using the ANOVA. * =p-value <0.05, ** = p-value <0.01, *** = p-value <0.001**** = p-value <0.0001.

**Figure 8:**
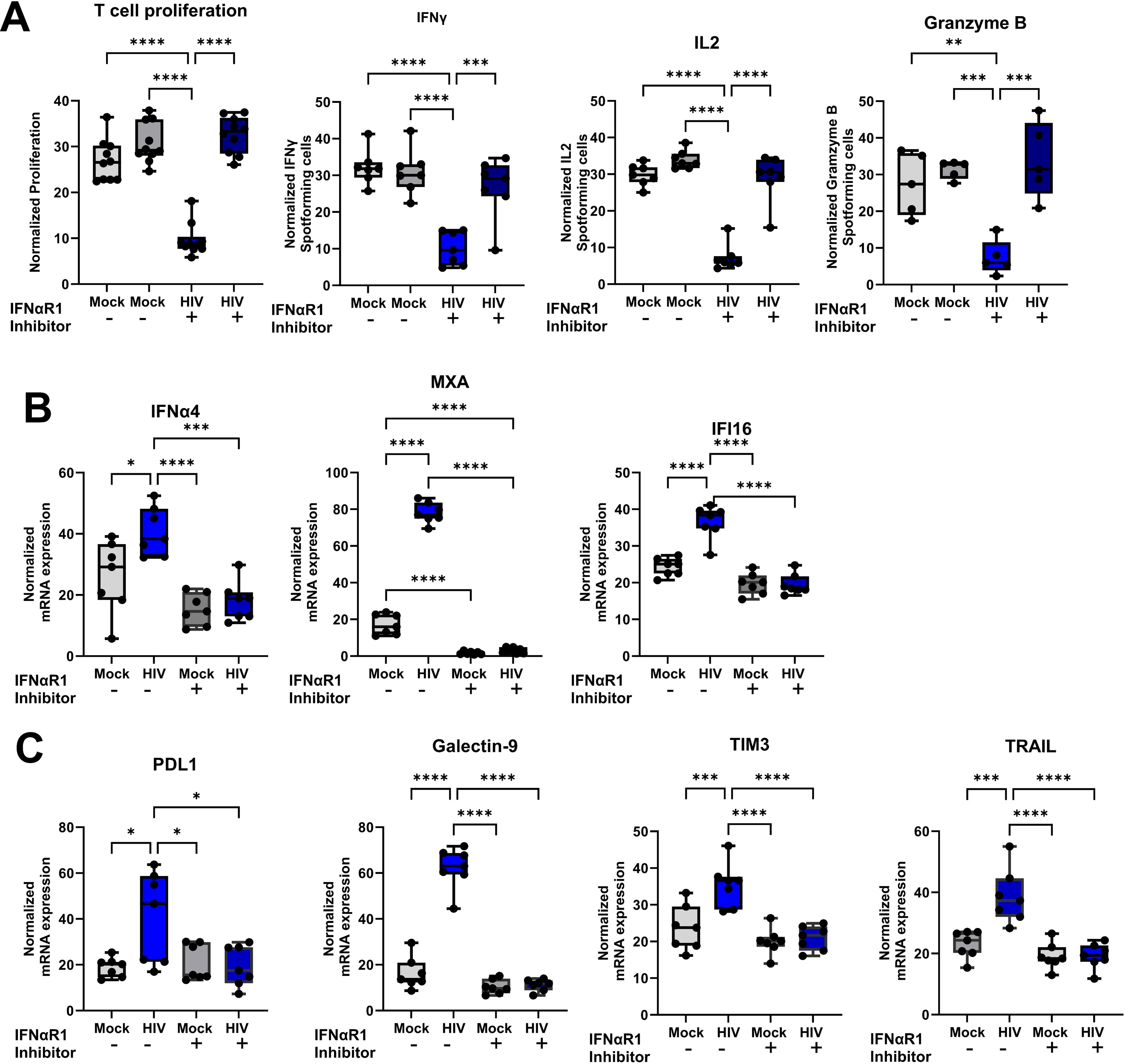
Targeting IFNaR1 by Anifrolumab restored immune function in HIV-induced suppressed DCs and T cells. Mature DC were exposed to HIV BaL and incubated for 24h. The untreated (mock) and HIV exposed DCs were harvested and exposed to 20ug/ml Anifrolumab for 30 min before the DCs were added to naïve T cells and co-cultured (1:10). The DC-T cell co-culture was re-stimulated on day 7 with the same DCs as the initial stimulation. One day after re-stimulation, the T proliferation was assessed via H^3^ thymidine incorporation and IFNγ, IL2 and granzyme B producing T cells by ELISPOT **(A)**. The effects of Anifrolumab on mRNA transcripts levels for IFNα4, IFI16, and MXA, and IFI16 **(B)** and PD1L, Galactin9, TIM3, and TRAIL) **(C)** were assessed by qPCR. Statistical significance was determined using the ANOVA. * =p-value <0.05, ** = p-value <0.01, *** = p-value <0.001**** = p-value <0.0001.

## Discussion

Here, we deciphered the HIV components and mechanisms underlying the immune suppression induced by HIV-exposed DCs when activating T cell responses and found major contribution of the cGAS/IFI16-STING signaling and type I IFN signaling pathways in the ensuing immune suppression.

Most individuals with untreated chronic HIV, which can be maintained during ART treatment, experience significant impairment in immune cell populations such as DCs, T cells, NK cells, B cells, and MAIT cells, often due to the sustained immune activation^13–15^. Immune activation is driven by multiple factors including prolonged elevated type I IFN levels and this is detrimental to the immune system function, which has been shown to contribute to disease progression in both HIV and simian immunodeficiency virus (SIV) infection^10,11^. Throughout HIV infection, HIV skews DC function, altering cytokine profiles, inducing partial DC maturation, and contributing to chronic immune activation and defective DC stimulation of T cells^13,14,35^. The observed dysfunction in T cells is in part attributed to the upregulation of coinhibitory molecules and loss of effector functions, further contributing to immune exhaustion. We have previously demonstrated that HIV-1 impairs T cell functionality through DC-mediated cell to cell contact with T cells^18,19^. DCs do not normally support a strong type I IFN after exposure to HIV, which utilizes the HIV restriction factors SAMHD1, and TREX1 to avoid cytosolic sensing by PRRs^36,37^. Nonetheless, our recent finding demonstrated a strong type I IFN profile in HIV-exposed DCs of DC-T cell co-cultures^20^, clearly indicating altered cellular programming supporting type I IFN responses. The prolonged type I IFN response observed in a setting of DC-T cell crosstalk in the presence of HIV may result from enhanced expression of the STAT1/STAT2/IRF9 transcription complex, sustaining ISG transcription^38^. Alternatively, it could stem from impaired IFNAR recycling, leading to receptor accumulation at the cell surface or increased endosomal uptake necessary for JAK-STAT signaling activation^39^. Dysregulation of downstream signaling, such as reduced ISG15 expression, may also contribute by destabilizing USP18, a negative regulator that inhibits IFNAR assembly and type I IFN signaling^38^.

Both the late-stage lab adapted HIV-1 BaL and the transmitted/founder HIV-1 THRO gave rise to suppressed T cell proliferation and impaired functionality and similar levels of type I IFNs responses, suggesting a conserved mechanism of immune suppression between different HIV-1 strains, and that HIV BaL is a good model virus to study HIV induced immune suppression.

The HIV replication cycle involves an array of nucleotides/replication intermediates, such as RNA genomes, ssDNA and dsDNA that are recognized and targeted by the host intracellular defense including RNA and DNA sensors, which is counteracted by HIV to avoid recognition ^40^. Our study reveals that HIV-1-derived oligonucleotides, i.e., ssDNA, dsDNA, and RNA:DNA hybrid transfected into the cytosol of DCs participated in the suppressed T cell proliferation, where ssDNA had a key role. The ssDNA, during HIV-1 replication, folds into structures resembling dsDNA which are recognized by the STING pathway ^23^. The results from our study suggest that virus derived oligonucleotides modify DCs and play a crucial role in HIV-1-induced immune suppression. During HIV infection the viral genome is protected by the viral capsid, protecting it from degradation and recognition by cytosolic sensors. If the capsid is disrupted, that leads to enhanced sensing of virus derived oligonucleotides by e.g., cGAS^41–43^. In our system it might be the creation of an environment that disrupts the viral capsid, exposing the viral genome that is the reason for the prolonged and strong type I IFN response we detect.

The mRNA profile sequenced in DCs derived from the HIV exposed DC-T cell co-culture highlighted the significant role of intracellular sensors such as cGAS and STING, and type I IFN pathways in the HIV-1 induced suppression of DCs. This is in accordance with other studies where cytosolic RNA and DNA sensors such as cGAS, RIG-I like receptors, and IFI16 have been implicated in recognizing HIV-1 and activating STING signaling in HIV infected ^23,25,44^. We found that addition of exogenous IFNα2 to the DC-T cell co-culture at concentrations detected in HIV exposed co-cultures, induced similar suppressed T cell responses as HIV exposure, suggesting that type I IFNs play a direct role in mediating the induction of impaired immune responses as seen in vivo in HIV and SIV infections^11,45^.

Our data showed that cytosolic HIV ssDNA and dsDNA mirroring the effects observed for HIV-1, which suggests that replication intermediates/nucleotides play a crucial role in HIV-1-induced immune suppression. The dsDNA-like structure is recognized as cytosolic DNA by the IFI16-STING pathway and activates type I IFN secretion^46^ and this has been shown in cells such as THP1, and monocytes^46^ and now by us in DCs, underscoring the unique immunosuppressive properties of structured ssDNA. Notably, targeting STING via intracellular cGAMP and agonists suppressed T cell function and induced IFNβ, MXA, and PDL1 expression, reinforcing the involvement of STING signaling cascade and subsequent type I IFN in the reprogramming of DCs to induce immune impairment in our co-culture system and in individuals with chronic HIV infection^47^. HIV-1 has evolved several escape mechanisms to counteract the STING-induced type I IFNs, e.g., by glutamylation of HIV p6 that inhibits STING and TRIM32 interactions, and by HIV Vif interaction with SHP-1 that leads to STING inhibition^48,49^. Hence, there is ‘tug-of-war’ between HIV blocking the STING pathway and the subsequent activation induced by its oligonucleotides produced during the viral life cycle.

Our findings suggest that RIG-I signaling had no significant role in HIV-induced immune suppression. This aligns with previous findings demonstrating that HIV-1 evades RIG-I recognition through mechanisms such as viral RNA modifications and sequestration of RIG-I signaling components^50^. The HIV engagement of both cGAS and RIG-I simultaneously has been indicated in enabling advanced innate recognition of HIV-1 by myeloid DCs in elite controllers ^51^, but if this occurs in our system needs to be further elucidated.

IFI16 is an important regulator to control HIV replication, independent of immune sensing via inhibiting the transcription factor SP1 in the nucleus of CD4+ T cells. This regulator function is especially efficient for non-clade C HIV-1 strains as these strains are in general more resistant to the IFI16 pathway^52^. Cytosolic DNA sensors such as IFI16 have been implicated in the recognition of HIV-1 ssDNA in THP1 cells and the subsequent activation of STING signaling^23^. Cells that lack IFI16 have uncontrolled replication, which in turn leads either to exhaustion of the cell or cell death^23^. We found that targeting the IFI16 in the DCs directly with an agonist lead to elevated type IFN and suppressed T cell proliferation to levels comparable to those observed with HIV-1. These results align with previous studies demonstrating that IFI16 plays a key role in sensing viral DNA^53,54^. Surprisingly, despite a similar T cell suppression, and type I IFN response pattern as HIV, activation of STING via IFI16 did not enhance the expression of co-inhibitory molecules or ISGs. IFI16-STING signaling seems to be a key component of the immune modulation seen in our system, but HIV-1 exposure clearly employs additional mechanisms that give rise to elevated expression of co-inhibitory molecules and further impair immune function that is missing when targeting only IFI16. The IFI16 sensor is interlinked with the cGAS-STING pathway in macrophages, by supporting cGAMP production and its downstream signaling, i.e., recruitment and activation of TBK1 ^55^, which might explain that individually targeting of cGAS or IFI16 sensor might not give the full effect that HIV induce in the DCs. High viral load drives the elevated expressions of IFI16 and cGAMP in HIV positive individuals. In addition, there was a correlation between CD38 and IFI16 in these patients, indicating that the DNA activation of IFI16 might be part of chronic immune activation^44^. Our data from the DC-T cell co-culture support this with high immune activation of type I IFN responses, and IFI16 leading to suppressed responses in the HIV-exposed setting. In chronic HIV infected individuals ART treated or untreated, there are impaired ISGs responses to DNA stimuli^44^, which could depend on the cell status due to the chronic activation found in these patients.

The elevated ISGs clearly indicated autocrine and signaling via IFNα/β receptors as part of the HIV induced immune activation leading to the suppressed responses. Prolonged type I IFN exposure and The IFNα/β receptors signaling via JAK/STAT/IRF9 has been linked to upregulation of inhibitory molecules, suppression of T cell function, and impaired antigen-presenting cell activity^56^. Blocking the IFNα, and IFNβ signaling via IFNAR using Anifrolumab fully restored the suppressive effects induced by HIV-1, including T cell proliferation, DC and T cell phenotypes and T cell functionality. This complete recovery rendered by Anifrolumab is a promising strategy to restore the immune function in people living with HIV and may also lead to lowered tissue viral DNA and improved general health as has been shown for ART treated SIV+ Macaques receiving anti-IFNα therapy using an anti-IFNα antibody^45^.

Overall, our results suggest that the IFI16/cGAS-STING pathway and subsequent type I IFN signaling with IFNα/β receptors contributes to HIV-induced immune suppression, but additional factors are likely involved in regulating the expression of co-inhibitory molecules. These findings reinforce the understanding that HIV exploits innate immune signaling to establish an immunosuppressive environment that hinders effective T cell responses and support the role of DC-driven immune regulation in HIV pathogenesis. Moreover, the restoration of immune function when targeting IFNαR via Anifrolumab, a drug already approved and on market for systemic lupus erythematosus and in phase 3 trials for lupus nephritis, systemic sclerosis, idiopathic inflammatory myopathies, and cutaneous lupus erythematosus, highlights the potential therapeutic value of targeting type I IFN signaling in HIV infection to alleviate immune dysfunction and improve DC and T cell functionality. Future studies should delineate how HIV-1 utilize the IFI16/cGAS-STING signaling and other factors involved in immune impairment and explore whether modulating the pathway can serve as a viable strategy to counteract immune suppression in HIV infection, both in untreated and ART treated individuals.

## Acknowledgment

AstraZeneca for the kind gift of Anifrolumab.

