## supplementary figures for "HIV-1 derived replication intermediates/oligonucleotides induce a type I IFN-dependent immune suppression via STING activation that can be restored by targeting IFNARI"

### Supplementary Figures and Tables Svanberg et al

**Supplementary table 1. Oligonucleotide Sequences.**

| Oligonucleotide | Sequence |
|---|---|
| ssDNA1 100 bases | GTC TCT CTG GTT AGA CCA GAT CTG AGG CAG CCT CAG ATG GCT AAC TAG GGA GAC CAC TGC TTA AGC CTC AAT ACA GCT TGT ATT GAG GCT TCA AGT AGT G |
| open form of ssDNA1 100 bases | CGT CTG GTT ACA TTA GAT CTG AGT CTG TGA GCT CTC TGG CTA ACT AGG GAA CCC ACT GAT AAT CGC TCA ATA AAG CTT GCC TTG AGT GCT TCA AGT AGT G |
| closed form of ssDNA1 100 bases | GTC TCT CTG GTT AGA CCA GAT CTG AGA GCA GCT CTC AGA TGG CTA ACT AGG GAG ACC ACT GCTT AA GCC TCA ATA CAG CTT GTA TTG AGT GCT TCA AGT AGT G |
| dsDNA forward | CCA TCA GA AAG AG GTT TAA TA TTT TTG TGA GAC CAT CGA AGA GAG AAA GAG ATA AAA CTT |
| dsDNA reverse | AAG TTT TAT CTC TTT CTC TCT TCG ATG GTC TCA CAA AAA TAT TAA ACC TCT TTC TGA TGG |
| Many loops | GTCTCTCTCCATAGACCAGATCTGAGCCTGGGAGCTCTCTGGCTAACTAG GGAACCCACTGCTTAAGCCTCAATAAAGCTTGCCTTGAGTGCA ACAAGTAGTG |
| DNA/RNA (sense) | ACATTAGTAGAACAAAATGGAATAACACTTTAAATCAAATAGCTACAAAA |
| DNA (antisense) | TTTTGTAGCTATTTGATTTAAAGTGTTATTCCATTTTGTTCTACTAATGT |
| RNA | ACA UUA GUA GAA CAA AAU GGA AUA ACA CUU UAA AUC AAA UAG CUA CAA AA |

Sequences taken from Jakobsen et al 2013[1]. Custom oligonucleotides were ordered from Eurofins genomics.

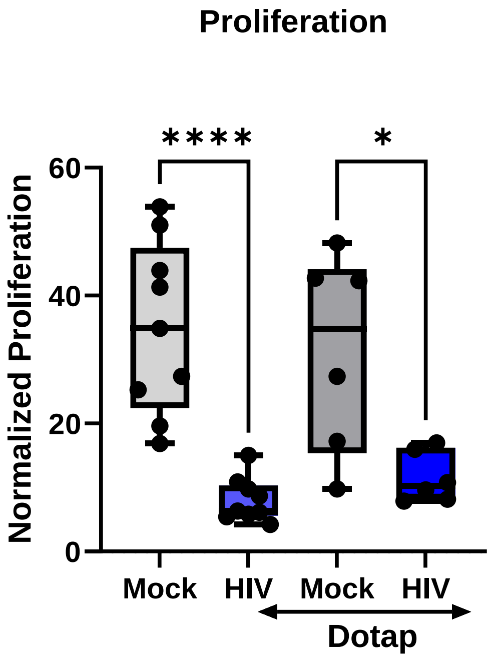


**Figure 1. DOTAP transfection had no effect on the induction of the HIV suppressive effects seen in the DC T cell coculture**

Mature DCs exposed to **mock or** HIV-1 BaL were untreated or treated by DOTAP transfection. The different DC groups were cocultured with naïve T cells (1:10) and the DC-T cell coculture was restimulated on day 7 with the same DCs as the initial stimulation. One day after restimulation, the T proliferation was assessed via $^{3}H$ thymidine incorporation **.** Statistical significance was determined using the ANOVA. * =p-value <0.05, ** = p-value <0.01, *** = p-value <0.00, 1**** = p-value <0.0001.
